# Using optogenetics to link myosin patterns to contractile cell behaviors during convergent extension

**DOI:** 10.1101/2021.02.25.432882

**Authors:** R. M. Herrera-Perez, C. Cupo, C. Allan, A. Lin, K. E. Kasza

## Abstract

Distinct spatiotemporal patterns of actomyosin contractility are often associated with particular epithelial tissue shape changes during development. For example, a planar polarized pattern of myosin II localization regulated by Rho1 signaling during *Drosophila* body axis elongation is thought to drive the cell behaviors that contribute to convergent extension. However, it is not well understood how specific aspects of a myosin localization pattern influence the multiple cell behaviors—including cell intercalation, cell shape changes, and apical cell area fluctuations—that simultaneously occur within a tissue during morphogenesis. Here, we use optogenetic activation (optoGEF) and deactivation (optoGAP) of Rho1 signaling to perturb the myosin pattern in the germband epithelium during *Drosophila* axis elongation and analyze the effects on contractile cell behaviors within the tissue. We find that uniform photoactivation of optoGEF or optoGAP is sufficient to rapidly override the endogenous myosin pattern, abolishing myosin planar polarity and reducing cell intercalation and convergent extension. However, these two perturbations have distinct effects on junctional and medial myosin localization, apical cell area fluctuations, and cell packings within the germband. Activation of Rho1 signaling in optoGEF embryos increases myosin accumulation in the medial-apical domain of germband cells, leading to increased amplitudes of apical cell area fluctuations. This enhanced contractility is translated into heterogeneous reductions in apical cell areas across the tissue, disrupting cellular packings within the germband. Conversely, inactivation of Rho1 signaling in optoGAP embryos decreases both medial and junctional myosin accumulation, leading to a dramatic reduction in cell area fluctuations. These results demonstrate that the level of Rho1 activity and the balance between junctional and medial myosin regulate apical cell area fluctuations and cellular packings in the germband, which have been proposed to influence the biophysics of cell rearrangements and tissue fluidity.

**STATEMENT OF SIGNIFICANCE:** Tissues are shaped by forces produced by dynamic patterns of actomyosin contractility. However, the mechanisms underlying these myosin patterns and their translation into cell behavior and tissue-level movements are not understood. Here, we show that optogenetic tools designed to control upstream regulators of myosin II can be used to rapidly manipulate myosin patterns and analyze the effects on cell behaviors during tissue morphogenesis. Combining optogenetics with live imaging in the developing fruit fly embryo, we show that acute perturbations to upstream myosin regulators are sufficient to rapidly perturb existing myosin patterns and alter cell movements and shapes during axis elongation, resulting in abnormalities in embryo shape. These results directly link myosin contractility patterns to cell behaviors that shape tissues, providing new insights into the mechanisms that generate functional tissues.

## INTRODUCTION

During development, tissues undergo dramatic changes in shape that are largely driven by patterns of contractile forces generated by the cellular actomyosin cytoskeleton (1–4). Patterns of myosin II localization and activity are responsible for producing spatially and temporally regulated cell behaviors that physically sculpt tissues and organs. In epithelial tissues, for example, planar polarized patterns of myosin localization at cell junctions as well as polarized flows of apical actomyosin are often associated with cell rearrangements that narrow and elongate tissues while pulsed, radial patterns of myosin at the apical surface of cells are often associated with apical constriction during tissue invagination (4, 5). Such myosin localization patterns are conserved, and significant insight into the roles of specific myosin patterns has been inferred from correlating myosin patterns with cell behaviors and tissue movements. However, gaining insight into how aspects of myosin localization and dynamics control distinct aspects of cell behavior requires experiments in which the myosin localization pattern is perturbed and the resulting cell behaviors analyzed.

Methods such as drug inhibition, genetic mutations, and protein knockdown or overexpression have been used to perturb myosin II and its regulators to study their functions during morphogenesis. These approaches can have significant limitations, depending on the process of interest. For example, during embryonic body axis elongation in *Drosophila*, drug inhibitors are typically injected into the embryo, limiting spatial control and potentially causing destructive effects at the injection site. Traditional molecular genetics perturbations cannot be easily or flexibly targeted at specific time points or to specific groups of cells at this stage of development in *Drosophila*, limiting spatial and temporal control. This makes it difficult to separate the effects of the perturbation on the tissue of interest from the effects in neighboring regions of the embryo or prior events during development. Thus, a major obstacle to understanding how patterns of actomyosin localization influence cell behaviors during tissue morphogenesis has been the lack of tools for flexible and precise manipulation of patterns of actomyosin contractility in vivo (6). Optogenetic technologies have emerged as powerful approaches for flexible and non-invasive manipulation of protein activities and cell behaviors (6–10). In particular, new optogenetic tools designed to manipulate actomyosin contractility with high spatiotemporal precision can now be used for studying the spatial and temporal requirements for myosin II in dynamic cell and tissue behaviors (11–20).

Convergent extension of the head-to-tail axis of the *Drosophila melanogaster* embryo is a powerful system for studying the mechanisms by which dynamic patterns of actomyosin contractility influence cell behaviors and tissue-level morphogenesis. During body axis elongation, the anterior-posterior (AP) axis of the embryo extends by two-fold. The majority of extension occurs during the first 30 min and is driven by cell intercalation (21–25) and cell shape changes (24, 26–28) in the germband epithelium. Just prior to the onset of body axis elongation, F-actin and myosin II become asymmetrically localized near adherens junctions at cell interfaces between anterior and posterior cell neighbors (“AP” or “vertical” edges), generating a planar polarized pattern of myosin II localization (22, 24, 29–31) and promoting cell intercalation (oriented cell rearrangement) that contributes to tissue elongation (22, 23, 32–35). In addition to junctional actomyosin, a dynamic actomyosin meshwork in the medial-apical domain of germband cells exhibits polarized flows and pulsatile behaviors that are thought to contribute to pulsatile fluctuations in cell junctions and apical cell areas and to cell intercalation (34–39). Recent theoretical and experimental studies point toward important roles both for active fluctuations in cell shape and junctions (40, 41) and for details of cellular packings within tissues (42–45) in the physics of cell rearrangements and the ability of a tissue to remodel and flow, i.e. tissue fluidity. It remains unclear exactly how cell intercalation, cell shape changes, and pulsatile cell area fluctuations are organized by the myosin localization pattern and together contribute to tissue shape change.

The patterns of myosin II localization during *Drosophila* axis elongation are regulated by the Rho/Rho-kinase signaling pathway (30, 35, 38, 46–50). In general, patterns of Rho activity are directed by Rho-specific guanine nucleotide exchange factors or RhoGEFs, which promote the active state of Rho, and by Rho GTPase-activating proteins RhoGAPs, which promote the inactive state of Rho (51–54). During axis elongation, the active state of the RhoGTPase *Drosophila* Rho1 is promoted by the exchange factors RhoGEF2 and Cysts/Dp114RhoGEF of the RhoGEF family (38, 46–50). Although RhoGAP activity is required for proper Rho signaling in some contexts (54– 56), RhoGAPs essential for *Drosophila* axis elongation have not yet been described. *Drosophila* RhoGAP71E has been shown to play a role, along with RhoGEF2, in organizing radial patterns of myosin activity that drive apical constriction during invagination of the presumptive mesoderm (56).

Because Rho activity controlled by RhoGEFs and RhoGAPs is thought to direct actomyosin contractility patterns in many contexts, it is an attractive target for optogenetic manipulation. Indeed, domains of RhoGEFs have been used successfully in optogenetic tools to activate actomyosin contractility and induce tissue folding in the early *Drosophila* embryo during cellularization and ventral furrow formation stages (16–18, 20), prior to the strong accumulation of myosin at adherens junctions present during axis elongation. Optogenetic tools based on RhoGAPs to inactivate Rho1 signaling and reduce actomyosin contractility have not yet been described to our knowledge. It remains unclear whether optogenetic technologies based on RhoGEFs and RhoGAPs will be sufficient to override and manipulate strong endogenous myosin patterns present in many tissues at later stages of development, for example during *Drosophila* axis elongation.

Here, we developed optogenetic tools to activate (optoGEF) or deactivate (optoGAP) Rho1 signaling. We used these tools to manipulate myosin patterns at the apical side of the germband epithelium during *Drosophila* body axis elongation and analyzed the effects on contractile cell behaviors. We show that uniform activation or inactivation of Rho1 signaling across the apical surface of the germband is sufficient to rapidly disrupt the endogenous planar polarized pattern of myosin at cell junctions on the timescale of 3-5 min, leading to distinct changes in junctional and medial myosin localization patterns in optoGEF and optoGAP embryos. These two perturbations to Rho1 activity both disrupt axis elongation and cell intercalation, but have distinct effects on cell area fluctuations and cell packings within the tissue that are linked with the changes in the medial versus junctional myosin pools. These studies demonstrate that acute optogenetic perturbations to Rho1 activity are sufficient to rapidly override the endogenous planar polarized myosin pattern in the germband during axis elongation. Moreover, our results reveal that the levels of Rho1 activity and the balance between medial and junctional myosin levels play key roles, not only in organizing the oriented cell rearrangements that are known to directly contribute to axis elongation, but also in regulating cell area fluctuations and cell packings within the tissue, which have been proposed to be important factors influencing the physics of tissue fluidity during development.

## RESULTS

### Optogenetic activation or inactivation of Rho1 signaling disrupts convergent extension in *Drosophila*

To dissect how patterns of actomyosin contractility influence cell behaviors that drive convergent extension, we used an optogenetic system to rapidly manipulate Rho1 signaling with high spatiotemporal precision at the apical surface of the epithelium of the developing *Drosophila* embryo. During convergent extension, myosin II in the germband epithelium is present in a planar polarized pattern at apical cell junctions (22, 23, 29) as well as in meshworks in the medial-apical domain of cells (34, 37) and is activated by the Rho1/Rho-kinase pathway (35, 38, 46–50). To manipulate patterns of Rho1 activity, we generated tools for blue-light gated recruitment of Rho1 regulators to the cell membrane using the CRY2/CIB1 heterodimerization system (7, 11, 12, 16) (Figure 1*A*).

**Figure 1.**
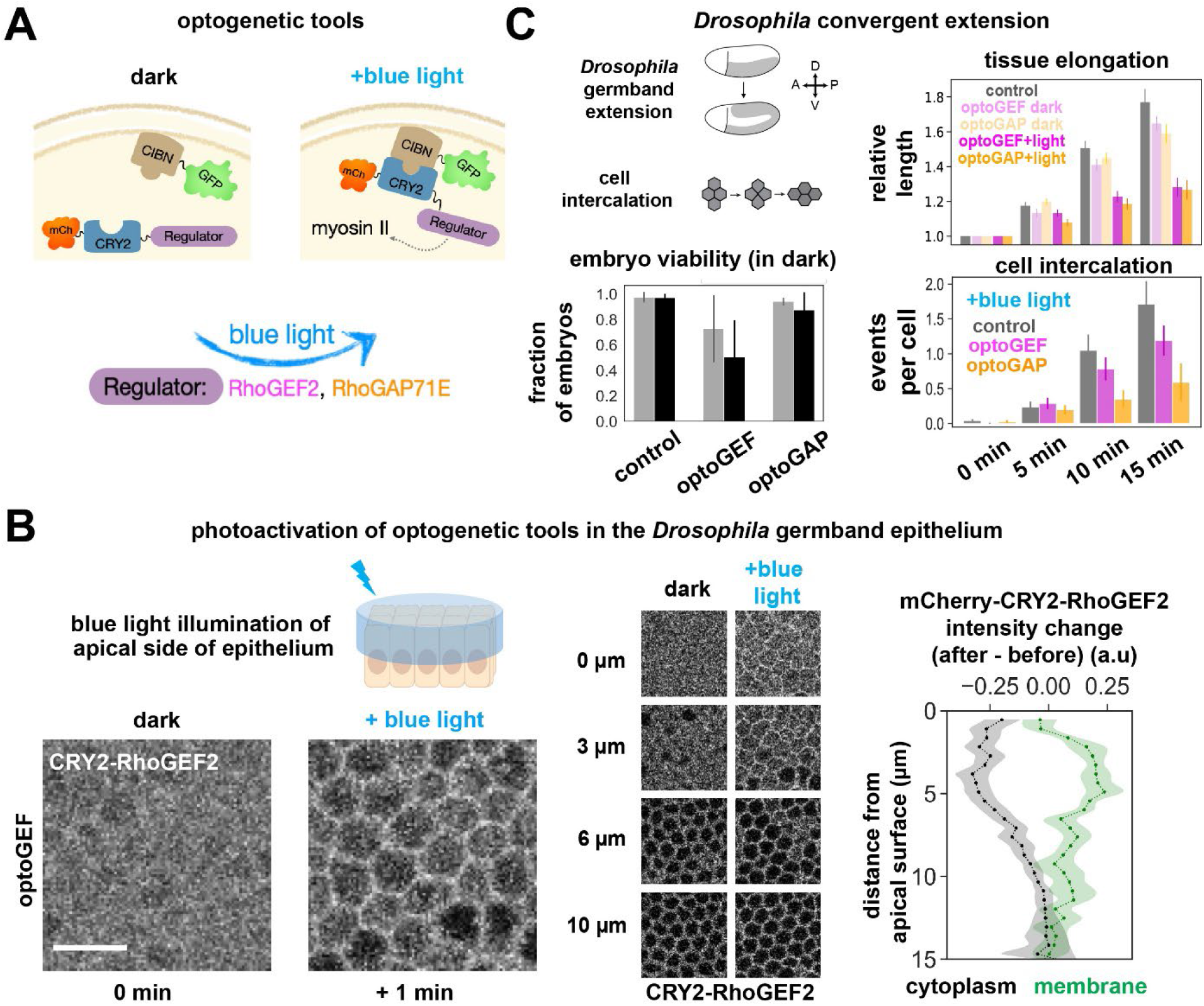
Optogenetic activation (optoGEF) or inactivation (optoGAP) of Rho1 signaling disrupts convergent extension during *Drosophila* axis elongation. (*A*) Schematic of optogenetic tools. CIBN is tagged with GFP and targeted to the cell membrane. The light-sensitive PHR domain of CRY2 is tagged with mCherry and fused to upstream myosin II regulators of the Rho/Rho-kinase pathway. Blue light illumination induces dimerization of CRY2 and CIBN, leading to accumulation of the regulator at the cell membrane. The full *Drosophila* RhoGEF2 (optoGEF) or RhoGAP71E (optoGAP) proteins were used to create the optogenetic tools. (*B*) Blue-light illumination of the apical surface of the germband epithelium in the *Drosophila* embryo to activate optogenetic tools during axis elongation. (*Left*) Prior to blue light exposure, the mCherry-CRY2-regulator component of the tool is localized in the cytoplasm of germband cells (optoGEF shown here). After 1 min of blue light exposure, mCherry-CRY2-regulator accumulates at the cell membrane and is depleted from the cytoplasm. Apical view, projection of z-slices from 2-8 µm. Bar, 10 μ m. (*Center*) Localization of the mCherry-CRY2-regulator component of the tool (optoGEF shown here) before and after blue light activation at different z-positions along the apical to basal axis of the epithelium: medial-apical (0 µm), junctional (3 µm) and lateral (6, 10 µm). (*Right*) Quantification of the change in intensity of mCherry-CRY2-RhoGEF2 at the cell membrane in optoGEF embryos after 1 min of blue-light activation. The greatest change is observed from 2-6 µm below the apical side of cells. Mean ± SEM between embryos is shown, n = 4 embryos, 6 cells per embryo. (*C*) (*Top left*) Schematics of convergent extension and cell intercalation during *Drosophila* body axis elongation. (*Bottom left*) Effects of expression of the optogenetic tools (in the absence of activating blue light) on the viability of embryos. Fraction of control, optoGEF, and optoGAP embryos that successfully completed axis elongation (gray) and hatch into larva (black). Mean ± SD between replicate experiments is shown, n = 163-227 embryos per group. (*Top right*) When the apical surface of the germband is illuminated with blue light during axis elongation, tissue elongation is reduced in optoGEF and optoGAP embryos compared to controls. Mean ± SEM between embryos is shown, n = 3-8 embryos per group. (*Bottom right*) Cell intercalation is reduced in optoGEF and optoGAP embryos compared to in controls. Mean ± SEM between embryos is shown, n = 3-5 embryos per group.

The first tool, optoGEF, was designed to control the cell membrane localization of *Drosophila* RhoGEF2, which promotes myosin activity in the germband and the presumptive mesoderm (38, 48, 56–61). We generated embryos that co-express CRY2-RhoGEF2 (the full RhoGEF2 fused to the light-sensitive CRY2 PHR domain, generated in this study) and CIBN-pmGFP (the N-terminal domain of the CRY2 binding partner CIB1 tagged with a cell membrane anchor and GFP) (7, 12). Blue-light induced accumulation of CRY2-RhoGEF2 at the cell membrane is predicted to increase Rho1 activity, leading to increased actomyosin contractility. The second tool, optoGAP, was designed to control the cell membrane localization of *Drosophila* RhoGAP71E, which plays a role in organizing radial patterns of myosin activity in presumptive mesoderm cells (56). The optoGAP tool (the full RhoGAP71E fused to the light-sensitive CRY2 PHR domain, generated in this study) is predicted to promote the inactive state of Rho1, leading to decreased actomyosin contractility.

To test the light-gated behavior of optoGEF and optoGAP during *Drosophila* axis elongation, we expressed the components of the tools in the early *Drosophila* embryo using the Gal4-UAS system. In the dark, the CRY2 component of the optogenetic tools localized to the cytoplasm (Figure 1*B*). We then photoactivated the apical surface of the epithelium with 488 nm laser light, scanned over the same volume used for imaging (Figure 1*B*). Blue light exposure led to rapid accumulation of the CRY2 component of the tools at the cell membrane and depletion from the cytoplasm (Figure 1*B*). The maximum membrane recruitment and cytoplasmic depletion occurred from 1-6 µm below the apical cell surface (Figure 1*B*), overlapping with the positions of myosin II at adherens junctions and in the medial-apical actomyosin meshwork, both of which contribute to contractile cell behaviors during convergent extension.

First, we analyzed the effects of the optogenetic tools on embryonic development in the absence of activating blue light. Embryos expressing the optoGAP tool displayed a slight reduction in embryonic viability, assessed by quantifying the fraction of embryos that go on to hatch into larva, compared to control embryos, while optoGEF embryos showed a greater reduction in viability (Figure 1*C*). We observed that some optoGEF embryos, those with very high expression levels of the tool components, displayed defects in egg shape, appearing more rounded and smaller than wild-type controls, or failed to cellularize. These results are consistent with the possibility that expression of the optogenetic tools at high levels can be deleterious in embryonic development, potentially due to over-expression effects of RhoGEF2 or RhoGAP71E as part of the transgenic optogenetic tools. We therefore restricted our attention to embryos expressing low to moderate levels of optoGEF or optoGAP in this study.

Next, we analyzed how photoactivation with 488 nm blue light influenced axis elongation in optoGEF and optoGAP embryos. Experiments were performed such that photoactivation of the germband was initiated at the onset of axis elongation at *t* = 0, which reduced potential effects on other developmental events in the embryo. Under continuous, uniform photoactivation by scanning 488 nm laser light over the apical surface of the germband during imaging, embryos expressing optoGEF elongated 1.28 ± 0.04-fold and embryos expressing optoGAP elongated 1.27 ± 0.06-fold during the first 15 min of axis elongation, significantly less than control embryos expressing only the CIBN-pmGFP component of the optogenetic system (1.77 ± 0.08-fold) (*P* = 0.018 and *P* = 0.019, respectively) or dark control optoGEF and optoGAP embryos not exposed to blue light (*P* = 0.065 and *P* = 0.066, respectively) (Figure 1*C*). Consistent with the reduced tissue-level elongation, the number of oriented cell rearrangements per cell was reduced to 1.2 ± 0.2 in optoGEF embryos (*P* = 0.28) and 0.67 ± 0.22 in optoGAP embryos (*P* = 0.03), compared to 1.7 ± 0.3 rearrangements per cell in control embryos (Figure 1*C,* *t* = 15 min). These results indicate that rapid optogenetic recruitment of Rho1 regulators to the apical surface of the germband epithelium is sufficient to disrupt axis elongation, providing an opportunity to dissect how Rho1 activity, and the patterns of actomyosin contractility it directs, influence contractile cell behaviors during convergent extension.

### Optogenetic control of Rho1 activity allows rapid manipulation of myosin patterns in the germband during axis elongation

Patterns of myosin localization and activity, and the resulting contractile forces generated, drive a wide array of cell behaviors that contribute to epithelial morphogenesis. For example, polarized actomyosin contractility, involving both the junctional and medial-apical pools of myosin, is required for cell intercalation during axis elongation (22, 23, 27, 29, 34, 48). While significant insight has been gained by correlating myosin patterns with cell and tissue behaviors during development, understanding how myosin patterns influence cell behaviors requires a direct perturbation approach. Drug injections do not allow fine spatial control and can be destructive to the embryo, and traditional genetic manipulations cannot be easily or flexibly targeted to specific tissues or developmental events at this stage of *Drosophila* development. To test if optogenetic manipulation of Rho1 activity in the germband might overcome the limitations of traditional perturbation approaches, we analyzed myosin localization patterns in optoGEF and optoGAP embryos.

To analyze how recruitment of Rho1 regulators at the apical surface of the tissue influences the spatial and temporal dynamics of myosin, we performed time-lapse confocal imaging in optoGEF and optoGAP embryos expressing an mCherry-tagged myosin regulatory light chain (*sqh*) transgene. In the absence of activating blue light, myosin II was present in a planar polarized pattern at cell junctions and in the medial-apical cortex in optoGEF and optoGAP embryos, similar to in control embryos (Figure 2*A-C*). We next tested how blue light illumination, beginning at the onset of axis elongation (*t* = 0) and continuing throughout the process, affected myosin localization. In control embryos expressing only the CIBN-pmGFP component of the system, myosin II was present at cell junctions and the medial-apical cortex of germband cells (Figure 2*D-D’’*). Just prior to the onset of axis elongation and before blue-light exposure, myosin began to accumulate more strongly at junctions between anterior and posterior cell neighbors (AP edges) than at junctions between dorsal and ventral cell neighbors (DV edges) (Figure 2*D*). During axis elongation and blue light exposure, myosin continued to accumulate at AP edges, resulting in a planar polarized pattern (Figure 2*D’-D’’*), consistent with previous studies of myosin planar polarity in wild-type embryos (22–24, 29–33).

**Figure 2.**
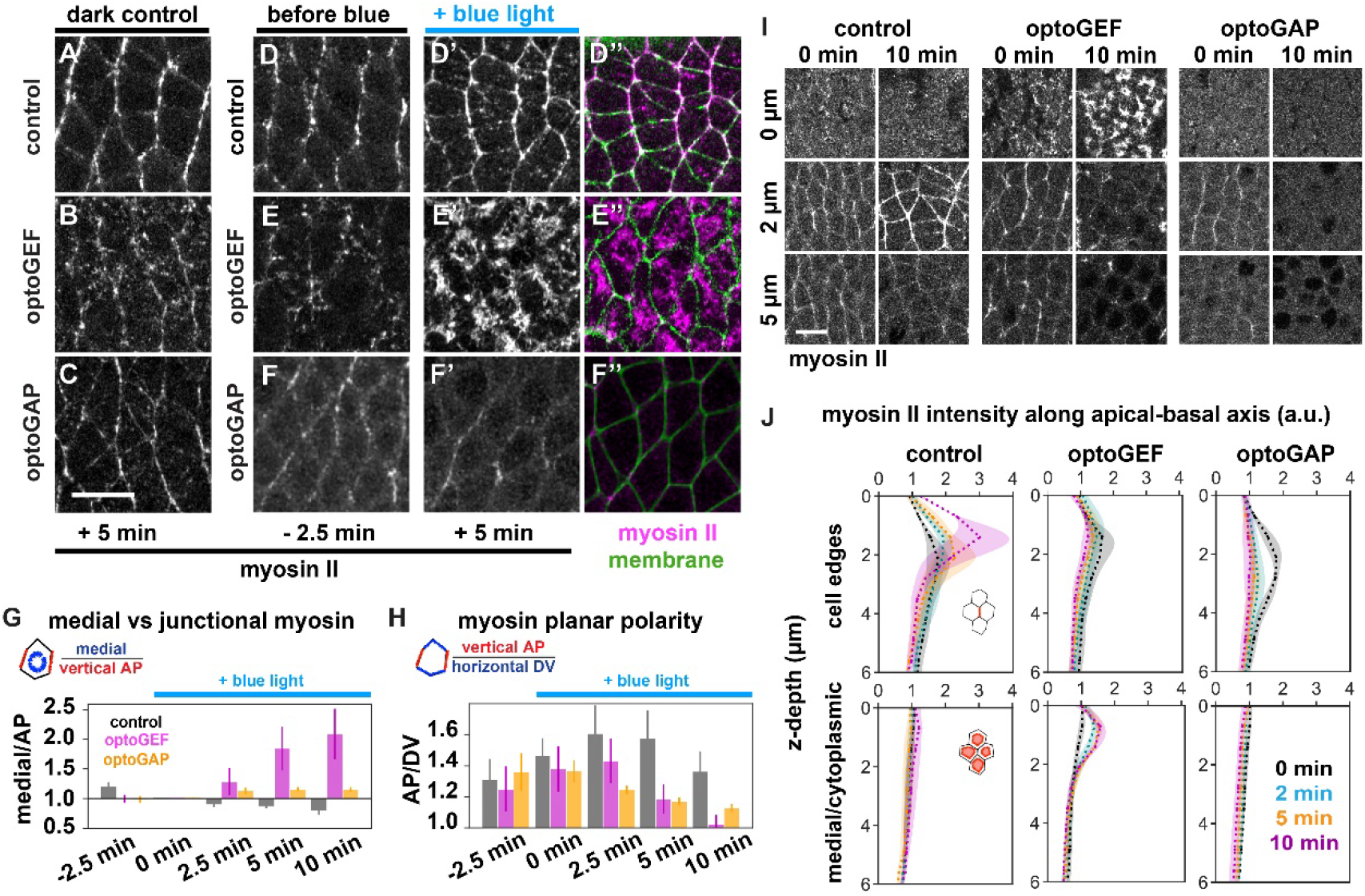
Optogenetic manipulation of Rho1 signaling with optoGEF or optoGAP alters junctional and medial myosin II localization patterns and disrupts myosin planar polarity during axis elongation. (*A-F*) Stills from movies of epithelial germband cells during *Drosophila* body axis elongation showing myosin II localization at the apical side of cells. Maximum intensity projection of z-slices in the most apical 5 µm. Myosin II (myosin) was visualized using an mCherry-tagged myosin regulatory light chain (*sqh*) transgene. (*A-C*) Dark control embryos without blue light activation (imaging only with 561 nm light). Myosin II is present in a planar polarized pattern, showing increased accumulation at “vertical” AP cell interfaces. (*D-F’’*) Embryos with 488 nm blue light activation (imaging with 488 nm and 561 nm light). (*D-F*) Two min prior to blue light exposure. (*D’-F’* and *D’’*-*F’’*) After 5 min of activating blue light exposure at the apical surface of the germband, optoGEF embryos showed an overall increase in cortical myosin levels (*E’-E’’*), and optoGAP embryos showed an overall decrease in cortical myosin levels (*F’-F’’*) compared to control embryos (*D’-D*’’). (*D’’*-*F’’*) Myosin II (magenta), cell membrane (green). Anterior, left. Ventral, down. Bar, 10 µm. (*G*) Ratio of mean myosin intensities in the medial-apical domain compared to at vertical AP junctions. Medial myosin accumulation relative to at AP junctions was increased in optoGEF compared to in control or optoGAP embryos: 2.09 ± 0.43 in optoGEF (*P* = 0.01), 1.16 ± 0.05 in optoGAP (*P* = 0.26), compared to 0.79 ± 0.07 in control embryos at *t* = 10 min. (*H*) Ratio of mean myosin intensities at vertical AP junctions compared to horizontal DV junctions. Myosin planar polarity was disrupted in optoGEF and optoGAP embryos (1.04 ± 0.06 in optoGEF and 1.13 ± 0.02 in optoGAP, compared to 1.41 ± 0.12 in control embryos at *t* = 10 min, *P* = 0.04 and *P* = 0.1 respectively). (*G,H*) Ratios of mean medial to mean AP or ratios of mean AP to mean DV intensities for each embryo at each time point were calculated. Mean ± SEM between embryos is shown, n = 3-4 embryos per condition. (*I*) Myosin localization at different z-planes along the apical-basal axis at 0 and 10 min of blue-light illumination, beginning from the onset of axis elongation. The most apical side of cells corresponds to z = 0. Bar, 10 µm. (*J*) Myosin intensity profiles along the apical-basal axis at AP cell edges (*top*), which represent contacts between anterior and posterior cell neighbors, and in the medial/cytoplasmic domain (*bottom*) at 0, 2, 5, and 10 min of blue-light illumination, beginning from the onset of axis elongation. Mean ± SEM between embryos is shown, n = 3 embryos per condition,10 cells per embryo.

In contrast, myosin localization patterns changed dramatically in cells of optoGEF embryos immediately following blue light exposure, indicating that these changes in myosin localization were induced by cell membrane accumulation of the Rho1 regulator RhoGEF2. In optoGEF embryos, myosin was initially localized at cell junctions prior to blue light exposure (Figure 2*E*). This myosin pattern was similar to that of control embryos (Figure 2*D*), but with slightly reduced junctional levels. Following blue light exposure starting at *t* = 0 to recruit RhoGEF2 to the cell membrane, myosin rapidly accumulated at the medial-apical side of cells. The rapid increase in medial myosin in optoGEF embryos was accompanied by a decrease in junctional myosin levels (Figure 2*E’*,*E’*’), producing a strong increase in the ratio of medial to junctional myosin and decrease in myosin planar polarity within 3-5 min of blue-light illumination. At *t* = 10 min, the ratio of medial to junctional myosin increased to a value of 2.09 ± 0.43 (*P* = 0.01) (Figure 2*G*) and myosin planar polarity decreased to a value of 1.04 ± 0.06 (*P* = 0.04) (Figure 2*H*). The resulting myosin pattern is reminiscent of the radial pattern in apically constricting cells in the presumptive mesoderm of *Drosophila* embryos (62–64). These results demonstrate that acute activation of optoGEF across the apical side of the germband rapidly disrupts the planar polarized pattern of junctional myosin localization organized by endogenous Rho1 signaling.

In optoGAP embryos, cortical myosin localization decreased following blue light exposure, indicating that these changes in myosin localization were induced by cell membrane accumulation of the Rho1 regulator RhoGAP71E. In these embryos, we observed gradual decreases in both medial and junctional myosin (Figure 2*F’*,*F’’*). This decrease in the cortical localization of myosin is consistent with the role of RhoGAPs in promoting the inactive state of Rho1. The changes in cortical myosin localization in optoGAP embryos were associated with a decrease in myosin planar polarity (1.13 ± 0.02 compared to 1.41 ± 0.12 in control embryos at *t* = 10 min, *P* = 0.1) (Figure 2*H*). These results demonstrate that acute activation of optoGAP across the apical side of the germband disrupts the levels of myosin at the cortex, including the planar polarized junctional pattern.

To test how the optogenetic perturbations influence myosin localization along the apical and lateral sides of cells, we analyzed myosin localization patterns at different z-positions in the confocal time-lapse movies. In control embryos, myosin became strongly enriched at cell contacts between anterior and posterior cell neighbors from 1.5-3 μ m below the apical side of cells (Figure 2*I,J*), consistent with localization at adherens junctions. Myosin was also present in the medial-apical domain at the very apical side of the cells from 0-1 μ m, and was not detectable at high levels along the lateral sides of cells below junctions (Figure 2 *I,J*). In optoGEF embryos, myosin became strongly enriched at the medial-apical side of cells and showed decreased junctional enrichment compared to controls, again with little detectable lateral accumulation (Figure 2 *I,J*). In contrast, myosin localization was reduced both at cell edges and in the medial-apical domain of optoGAP embryos (Figure 2 *I,J*).

Taken together, these findings demonstrate that activation of either optoGEF or optoGAP at the apical side of the germband, beginning at the onset of axis elongation, is sufficient to override the planar polarized myosin patterns that are organized by endogenous Rho1 signaling in the germband, converting the planar polarized junctional myosin pattern into a more medial-apical radial pattern in optoGEF or strongly reducing overall cortical myosin levels in optoGAP, over the timescale of a few minutes. In both optoGEF and optoGAP embryos, myosin planar polarity was significantly reduced, consistent with the strong reductions in cell rearrangements and tissue elongation in these embryos (Figure 1*C*). Such rapid, local, and non-invasive optogenetic perturbations provide a unique opportunity to study how myosin patterns affect contractile cell behavior during convergent extension.

### Optogenetic manipulation of Rho1 activity is associated with changes in apical cell area

The specific pattern of actomyosin localization and activity within cells is thought to drive a wide variety of cell behaviors. Within the germband during axis elongation, myosin displays a complex localization pattern; it is strongly enriched at cell junctions where it displays a planar polarized accumulation at vertical AP junctions (22, 23, 29) and it also present in a dynamic medial-apical meshwork that exhibits polarized flows toward AP junctions (27, 34, 35, 37, 48). While it is known that the planar polarized myosin pattern is required for oriented cell rearrangements that drive axis elongation (and our results of reduced myosin planar polarity, cell rearrangement, and tissue elongation in optoGEF and optoGAP embryos are consistent with this), it remains less clear how the balance between junctional and medial myosin influence other aspects of contractile cell behaviors in the germband, such as cell shape and apical cell area. To dissect how junctional and medial myosin localization patterns influence apical cell shape and area, we analyzed how optogenetically perturbed myosin localization patterns in the germband of optoGEF and optoGAP embryos influence these contractile cell behaviors.

First, we analyzed cell shapes during axis elongation, as cell stretching along the AP axis contributes to tissue-level elongation (24, 26, 28). In control embryos, the mean horizontal length of cells along the AP axis increased by 12 ± 4% at *t* = 10 min relative to at *t* = 0 min (Figure 3*A,B*), similar to previous studies of wild-type embryos (24, 27, 28, 45). Because cell stretching along the AP axis is thought to be associated with external forces from posterior midgut invagination and is also thought to depend on cell rearrangements in the germband that can relax this stretch (26, 28, 45, 65), we hypothesized that the reduced cell rearrangements in optoGEF and optoGAP embryos might contribute to increased stretching. Alternatively, we wondered if the increased medial myosin accumulation in optoGEF embryos might provide contractile tensions resisting the external forces and reducing cell stretching. We found that cell length changes were reduced in photoactivated optoGEF embryos (*P* = 0.67) and not strongly affected in photoactivated optoGAP embryos (*P* = 0.87) compared to controls, but the differences were not statistically significant (Figure 3*A,B*). These results suggest that changes in medial and junctional myosin patterns have the potential to impact cell stretching during axis elongation, although the effects in these studies were not consistently strong.

**Figure 3.**
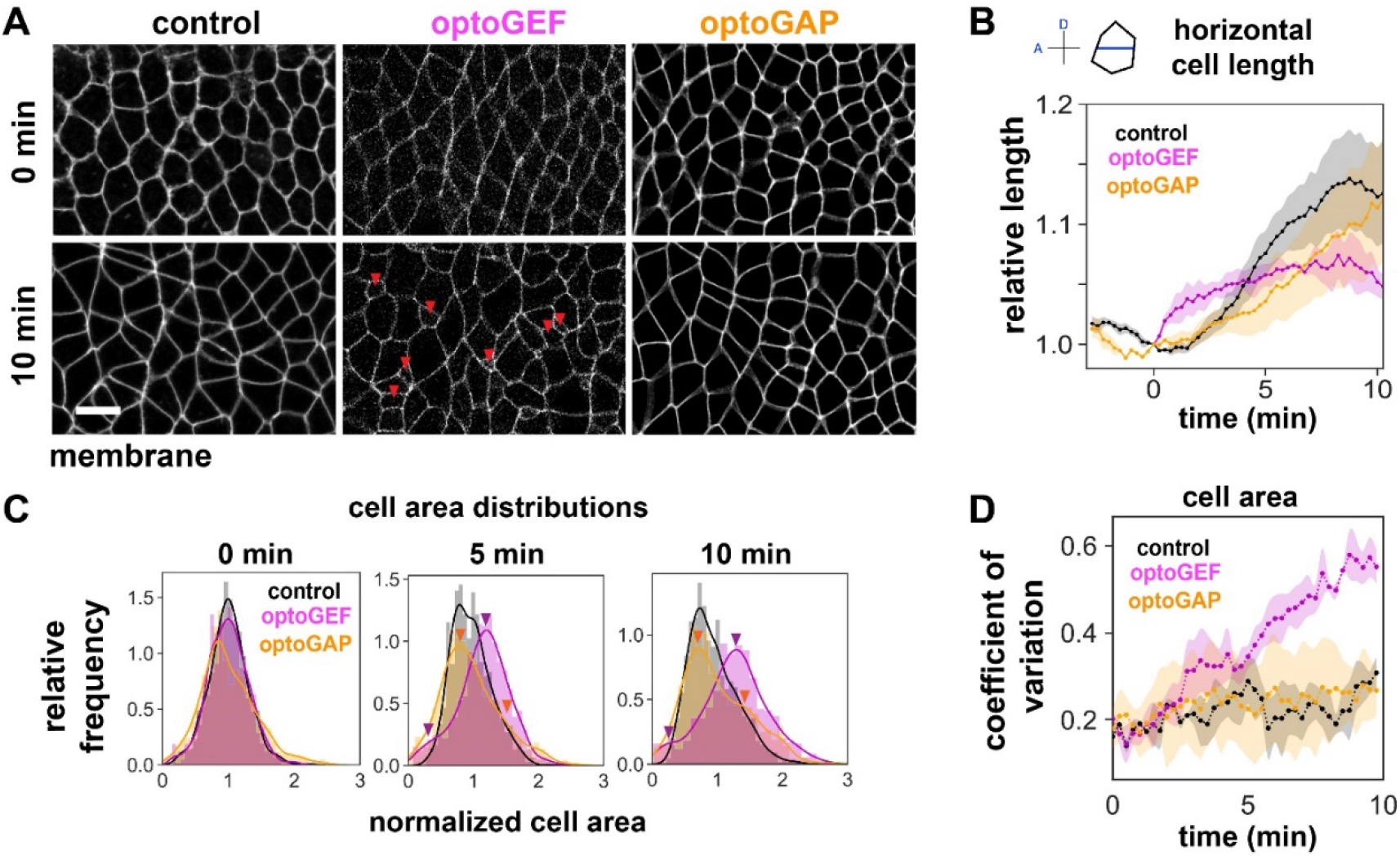
Perturbing Rho1 activity across the germband using optoGEF or optoGAP alters cell areas. (*A*) Stills from confocal movies of apical cell shapes in the germband during axis elongation. Some cells with small apical areas are observed in optoGEF embryos (red arrows). Anterior, left. Ventral, down. Bar, 10 μ m. (*B*) Mean cell length along the AP axis (horizontal length) normalized to the value at *t* = 0, a metric for changes in cell shape that contribute to tissue elongation. Mean ± SEM between embryos is shown, n = 3-5 embryos per condition. (*C*) Histograms of apical cell areas at 0, 5, and 10 min, normalized to the mean value at each point in each condition. The cell area distribution is unimodal in control embryos, but becomes bimodal in optoGEF and optoGAP embryos, indicating the presence of sub-populations with abnormally smaller or larger areas. (*D*) The coefficient of variation of apical cell area as a function of time in control, optoGEF, and optoGAP embryos. Cell area heterogeneity increases in optoGEF embryos compared to in control or optoGAP embryos. Mean ± SEM between embryos is shown, n=3-5 embryos per genotype.

Next, we analyzed cell areas in the germband tissue, as medial-apical actomyosin contractility plays key roles in apical constriction in a number of contexts, including ventral furrow invagination in the presumptive mesoderm of the *Drosophila* embryo. During ventral furrow invagination, a group of cells exhibiting strong medial-apical myosin contractility constrict their apical sides, leading to a significant apical cell area reduction and tissue invagination in this region (63). Remarkably, this apical cell constriction and invagination was reproduced via local optogenetic activation of Rho1 in small groups of cells in the early *Drosophila* embryo (16, 20). Naively, one might expect similar behavior here, with reduced apical cell area in the photoactivated germband of optoGEF embryos. However, when we activated Rho1 across the ventrolateral region of the germband in optoGEF embryos, we did not observe uniform apical cell area reductions across the tissue and instead observed the presence of a small fraction of cells with aberrantly small apical cell areas (Figure 3*A*, *red arrows*).

To quantify this heterogeneity, we measured the apical areas of cells within the tissue and plotted the data as a histogram at different time points during axis elongation. At the beginning of axis elongation (*t* = 0), control embryos display a Gaussian, unimodal distribution of cell areas (Figure 3*C*). Over time, the distribution widens somewhat as cells undergo cell rearrangements during axis elongation. In contrast, while optoGEF embryos start out with a unimodal cell area distribution similar to in controls at *t* = 0, we observe the appearance of a population of cells with drastic reductions in area while other cells display a modest increase in area at 5 min and 10 min of blue-light illumination, leading to a bimodal area distribution (Figure 3*C*). In optoGAP embryos, we observed a cell area distribution with some cells with somewhat increased area relative to the mean during axis elongation (Figure 3*C*).

To directly compare cell area heterogeneity in these different perturbations, we quantified the coefficient of cell area variation and plotted this as a function of time (Figure 3*D*). In control embryos, the variation in area among cells started out at a low level of 0.16 at *t* = 0 and increased by 1.9 ± 0.16-fold during the first 10 min of axis elongation. Cell areas in optoGAP embryos showed less change in area variation over time (1.5 ± 0.07-fold change from *t* = 0 to 10 min), with a similar coefficient of variation as control embryos at *t* = 10 min (*P* = 0.27). In contrast, cells in optoGEF embryos showed a greater increase in cell area variation, starting around 3 min after tool activation, reaching a 2.81 ± 0.42-fold increase at *t* = 10 min (*P* = 0.03). These results are consistent with the notion that decreased medial and junctional myosin in optoGAP cells leads to apical relaxation, associated with relatively uniform apical cell areas across the germband. In contrast, the increased medial myosin in optoGEF cells would be consistent with increased apical contractility. This apical contractility is translated into cell area reductions in some cells and area increases in other cells, instead of uniform cell area decreases in the tissue. Taken together, these results demonstrate that rapid optogenetic perturbation of Rho1 activity alters the apical areas of epithelial cells within the germband.

### Cell area changes over time are linked to junctional and medial myosin levels

To further link myosin patterns to contractile cell behaviors and investigate the origins of the cell area heterogeneity in the germband of optoGEF embryos, we next investigated how myosin localization and dynamics influence cell area changes in single cells over time. Cells in the germband exhibit dynamic apical area fluctuations or “pulses” that correlate with pulsatile medial myosin dynamics and pulsatile changes in the length of shrinking cell edges during cell intercalation (34, 37, 66). In one model, the cell population with aberrantly small areas in optoGEF embryos might represent cells that are progressively shrinking or ratcheting down their apical area over time. Alternatively, the small cell population might reflect cells that are fluctuating or pulsing, but not progressively changing apical area, and just happen to have a small area at the time point captured. To distinguish between these possibilities and directly link myosin pattern to contractile cell behavior at the single cell level, we tracked and quantified cell area, junctional myosin, and medial myosin in groups of individual cells over time in control, optoGEF, and optoGAP embryos (Figure 4).

**Figure 4.**
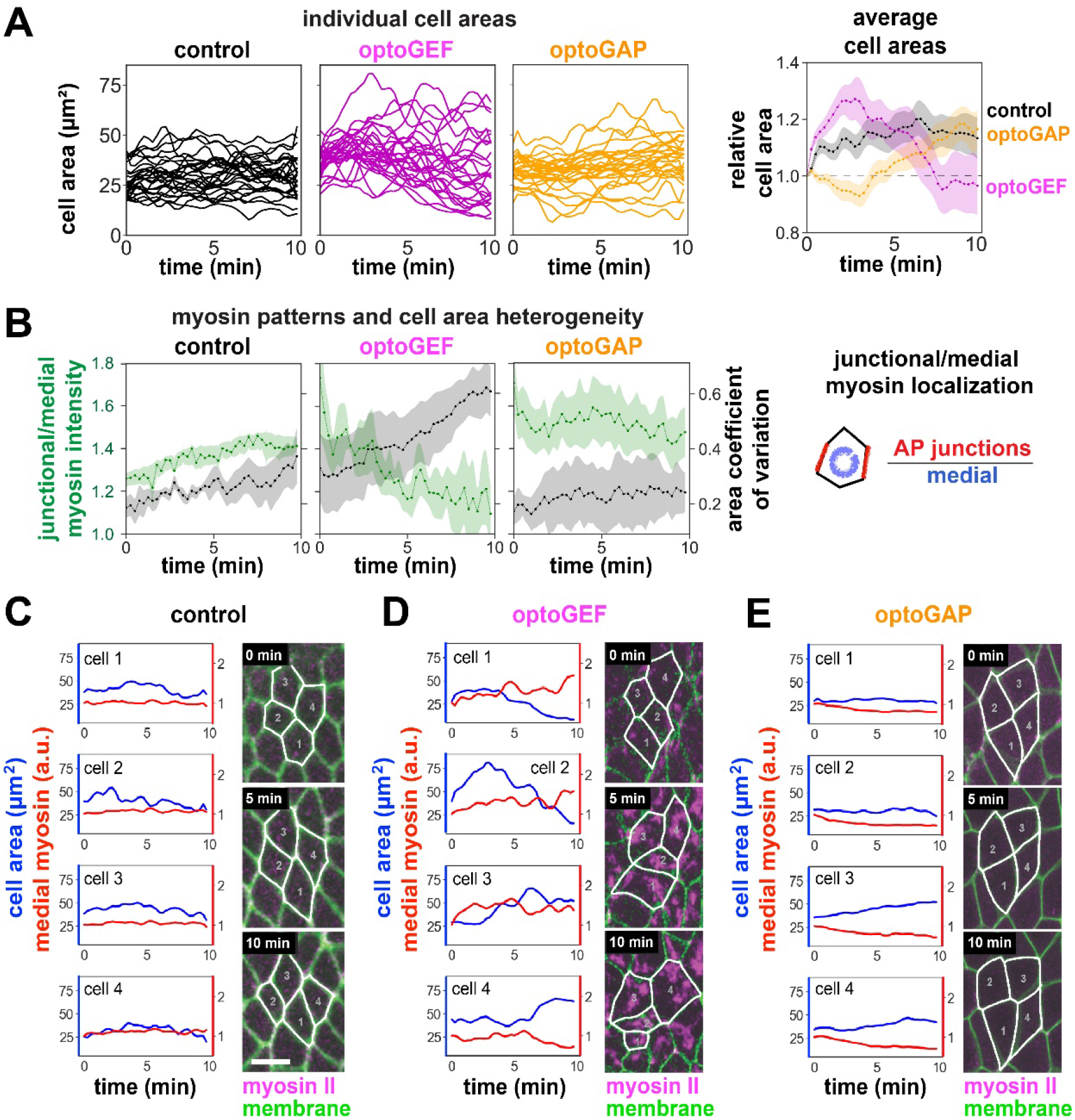
Changes in apical cell area are associated with changes in medial and junctional myosin patterns during convergent extension. (*A*) (*Left*) Apical cell area over time for individual germband cells in control, optoGEF, and optoGAP embryos during 10 min of blue-light illumination of the apical surface of the tissue during axis elongation. (*Right*) Average cell areas normalized to the values at *t* = 0. (*B*) Accumulation of myosin at vertical AP cell junctions relative to in the medial-apical domain (*green*) and coefficient of variation of apical cell area (*black*) in groups of cells analyzed over time. Maintenance of relatively high junctional myosin levels compared to medial myosin levels in control and optoGAP embryos is associated with low cell area heterogeneity. Decreases in junctional myosin and concomitant increases in medial myosin in optoGEF embryos are associated with increased cell area heterogeneity. Mean ± SEM between embryos is shown, n = 12 cells per embryo, 3-4 embryos per genotype. (*C-E*) Examples of apical cell area and medial myosin intensity in individual cells over time in control (*C*), optoGEF (*D*), and optoGAP (*E*) embryos. Cell area (*blue*) and medial myosin intensity (*red*) in four neighboring cells. Stills from confocal movies showing cells at 0, 5, and 10 min. Myosin II labeled with mCherry-tagged myosin regulatory light chain, magenta. Cell membrane labeled with CIBN-pmGFP, green. Overlaid cell outlines, white. Bars, 10 μ m.

There was heterogeneity in cell area changes over time between neighboring cells within a tissue in control embryos as well as in optoGEF and optoGAP embryos (Figure 4*A,C-E*). In control embryos, the areas of individual cells fluctuated over time but maintained some average value during axis elongation (Figure 4*A,C*). This is reflected in a relatively constant mean cell area in the tissue over time (Figure 4*A*). Similar behavior was observed in photoactivated optoGAP embryos, although the area fluctuations were less prominent (Figure 4*A,E*). In contrast, in photactivated optoGEF embryos, we observed that the pulsed cell area fluctuations were increased in magnitude and that some cells progressively decreased in area over time while neighboring cells progressively increased their apical area (Figure 4*A,D*). On average, this resulted in a net decrease in apical cell area and a high variation between cells in the optoGEF group compared to in controls at the 10 min time point. Interestingly, we find that the increase in cell area heterogeneity in optoGEF embryos correlates with the decrease in junctional myosin and increase in medial myosin following optogenetic perturbation (Figure 4*B*). In contrast, control and optoGAP embryos maintain low area heterogeneity, which correlates with maintaining high junctional relative to medial myosin levels, even when overall levels of myosin are decreased in optoGAP embryos (Figure 4*B*).

Taken together, these results suggest that maintaining a balance between junctional and medial myosin contributes to apical cell area maintenance across the tissue. The variability in cell areas across the photoactivated germband of optoGEF embryos that have increased medial myosin is distinct from the more uniform apical constriction in cells of the presumptive mesoderm during ventral furrow formation (63) and in groups of cells with optogenetically induced Rho1 activity in the early embryo (16, 20). One possible explanation is the larger size of the activated region used in this study. If all cells in a large region of the germband tissue try to contract at the same time, perhaps the tissue-scale deformation cannot be accommodated by surrounding tissue, leading to some activated cells contracting and others expanding. Other possibilities include cell-to-cell variability in expression of the optogenetic tools or changes to the actin cytoskeleton or cell-cell adhesions that contribute to apical cell area maintenance. Thus, optogenetic manipulation to activate Rho1 signaling across the germband, which is associated with increased medial and decreased junctional myosin, leads to greater net apical cell area changes and heterogeneous decreases in cell area. In contrast, inactivating Rho1 signaling across the germband, which is associated with decreased medial and junctional myosin, leads to less prominent cell area changes.

### The magnitudes of cell area fluctuations are tuned by optogenetic perturbation of Rho1 activity

Fluctuations or pulses in myosin accumulation and associated changes in apical cell shape have been described in many epithelia and are thought to play key roles in diverse cellular processes during morphogenesis, including apical cell constriction and cell intercalation (1, 67). Moreover, active fluctuations in cell shape and actomyosin accumulation have been proposed to play key roles in promoting cell rearrangement and tissue fluidity in some contexts (40, 41). While cell area fluctuations have been reported in the germband during axis elongation (34, 37, 66), their role in the process remains unclear. We wondered if the short timescale fluctuations in cell area that we observed in control and optoGEF embryos were correlated with myosin dynamics.

To test this, we first measured changes in cell area and in medial myosin between successive 15 s time points in confocal time-lapse movies (Figure 5*A-C*). In control embryos, the peaks in cell area change pulses had an amplitude of 5.0 ± 0.1 μ m^2^ and period of 2.59 ± 0.11 min (Figure 5*A,D,E*), comparable to in prior reports (37, 66). The amplitude of changes in apical cell area in optoGEF embryos was increased to 6.0 ± 0.2 μ m^2^ (*P =* 0.018). In contrast, large pulses were nearly absent in optoGAP embryos (Figure 5*A-C,D,E*). The time period between pulses was not significantly altered in optoGEF (2.49 ± 0.99 min, *P =* 0.66) or optoGAP (2.19 ± 0.07 min, *P =* 0.19) compared to in control embryos (Figure 5*E*).

**Figure 5.**
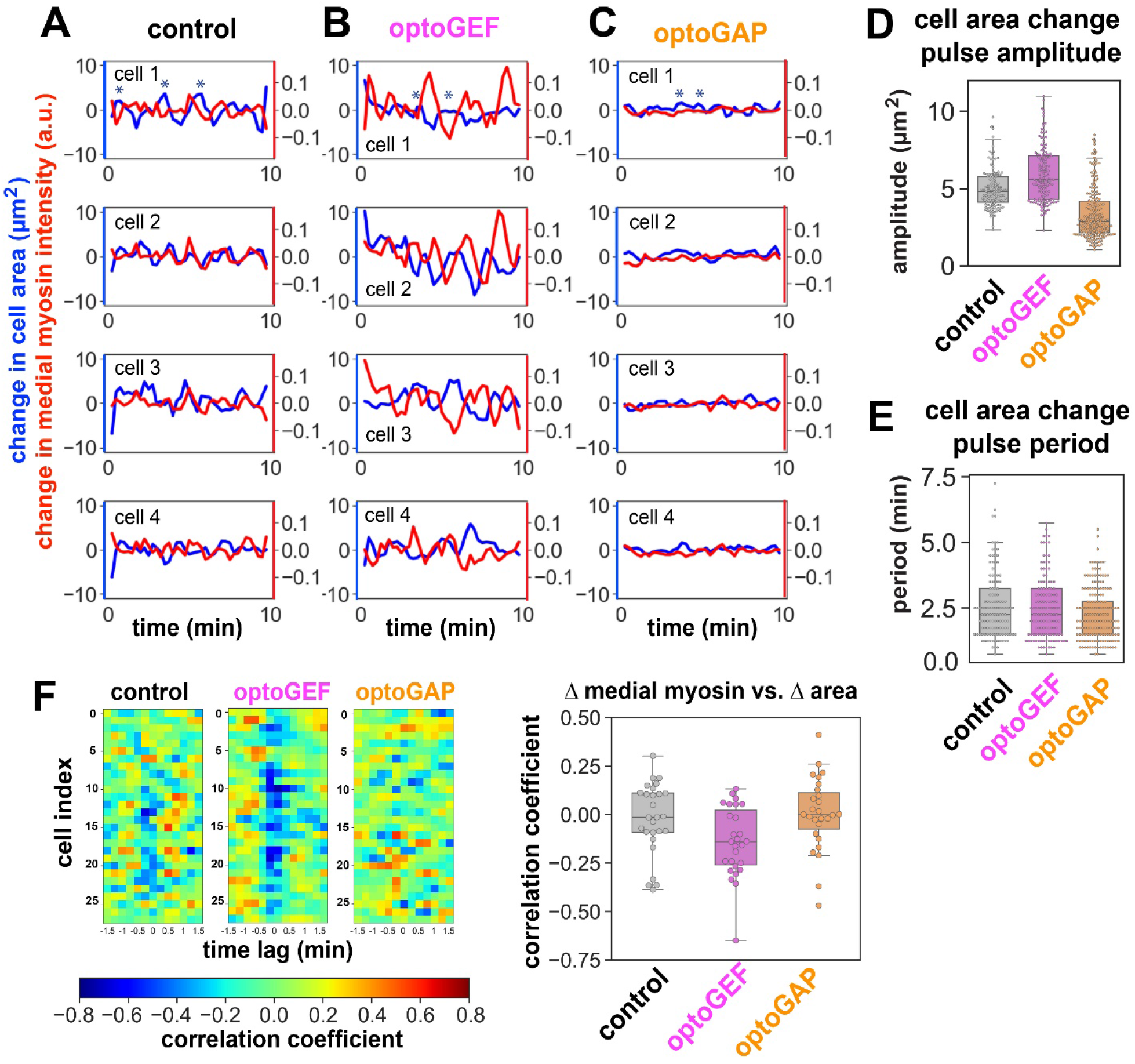
Apical cell area fluctuations are increased in optoGEF and decreased in optoGAP embryos. (*A-C*) Changes in apical cell area (*blue*) and changes in normalized medial myosin intensity (*red*) between 15 s time points in four typical cells in control (*A*), optoGEF (*B*), and optoGAP (*C*) embryos photoactivated with blue light. Example pulses indicated by * symbols. (*D*) Amplitude of area change pulses is increased in optoGEF and decreased in optoGAP compared to control embryos (optoGEF: *P =* 0.018, optoGAP: *P =* 6 × 10^−9^). The error in area associated with cell segmentation was 3 µm^2^, and so pulses less than 3 µm^2^ in amplitude were not included in analysis for control and optoGEF embryos. The majority of fluctuations in optoGAP embryos were less than 3 µm^2^ in amplitude, and so all peaks are shown in this dataset. Each point corresponds to one cell area pulse, n = 12 cells per embryo, 3-4 embryos per genotype. (*E*) Time period between successive cell area pulses (optoGEF: *P =* 0.66, optoGAP: *P =* 0.19 compared to controls). (*F*) Cross-correlation between changes in medial myosin and changes in apical cell area. (*Left*) Correlation coefficients at different time lags for individual cells. (*Right*) Correlation coefficient with zero time lag (optoGEF: *P =* 0.18, optoGAP: *P =* 0.83 compared to control embryos). n = 28 cells from 3-4 embryos per genotype.

Next, we tested whether the cell area fluctuations were correlated with the medial myosin fluctuations. We did not find a significant temporal correlation in control embryos over the 10 min period analyzed at the beginning of axis elongation (mean correlation coefficient *=* −0.019) (Figure 5*F*). This is in contrast to correlations between cell area and medial myosin changes observed at later stages of axis elongation, when medial myosin accumulation was found to precede cell area changes by several seconds (37, 66); this difference is potentially explained by the higher levels of medial myosin at the later stages of axis elongation. The correlations between cell area changes and medial myosin changes were modestly increased in optoGEF embryos (mean correlation coefficient *=* −0.14, *P =* 0.18) (Figure 5*F*). This is consistent with trends observed in previous studies in wild-type embryos later in axis elongation (37, 66) and in optogenetically-induced actomyosin contractility mimicking furrow formation (16) that report correlations between medial myosin accumulation and cell area changes. Taken together, these results demonstrate that cell area fluctuations are tuned by Rho1 activity in the germband, with the pulse magnitudes increased in optoGEF and decreased in optoGAP embryos.

## DISCUSSION

Patterns of actomyosin contractility influence multiple aspects of cell behavior within tissues that contribute to tissue-level morphogenesis. Here, we used optogenetic tools to activate or deactivate Rho1 signaling across the apical side of the germband epithelium during *Drosophila* axis elongation to manipulate myosin patterns and analyze the effects on cell behaviors within the tissue. We demonstrate the ability to rapidly override the endogenous planar polarized myosin pattern with both an optoGEF and an optoGAP tool. In optoGEF embryos, the planar polarized pattern of myosin localization at cell junctions is transformed into a radial pattern at the medial-apical cortex of cells following recruitment of RhoGEF2 to the apical cell membrane. In optoGAP embryos, recruitment of RhoGAP71E results in a decrease in myosin accumulation at the cell cortex in both the junctional and medial-apical domains, abolishing the endogenous myosin pattern. Both optogenetic perturbations resulted in a loss of myosin planar polarity and strong reductions in cell intercalation and tissue elongation. However, optoGEF and optoGAP had distinct effects on cell shapes and pulsatile cell area fluctuations, which can contribute to tissue behavior during convergent extension. While depletion of cortical myosin in optoGAP embryos was associated with a reduction in apical cell area fluctuations, medial myosin accumulation in optoGEF embryos led to enhanced apical area fluctuations and heterogeneous decreases in apical cell area.

For this study, we developed two CRY2/CIBN-based optogenetic tools to manipulate actomyosin in *Drosophila*: optoGEF to activate and optoGAP to deactivate Rho1 signaling. To our knowledge, a RhoGAP-based optogenetic tool to manipulate actomyosin contractility in *Drosophila* has not been previously described. Related optogenetic approaches based on RhoGEFs using the DHPH catalytic domains of RhoGEF2 in a CRY2/CIBN system (16–18) or the catalytic DH domain of LARG GEF in a LOV-domain-based system (20) have previously been used to manipulate Rho1 activity in the *Drosophila* embryo. The optoGEF tool described here is distinct from these prior tools in that it includes the full RhoGEF2 protein, instead of only the catalytic domains, and so has the potential to recapitulate additional aspects of RhoGEF2 function. Prior studies using RhoGEF-based optogenetic tools focused on manipulating Rho1 activity and actomyosin contractility during early stages of embryonic development, including during cellularization and ventral furrow formation (16–18, 20), revealing the potential to reconstitute aspects of morphogenesis via ectopic, local Rho1 activation in the epithelium prior to the appearance of strong myosin localization patterns at adherens junctions. In contrast, in this study we perturb myosin localization and activity in the epithelium at a later stage of development when there is an existing planar polarized pattern of myosin localization, and we analyze how these perturbations influence cell behaviors during axis elongation. Consistent with prior observations (16, 20), we find that optogenetic activation of Rho1 on the apical side of the *Drosophila* epithelium leads to rapid accumulation of myosin in the medial-apical domain of cells. We observe pulsatile apical cell area fluctuations in optoGEF embryos, consistent with pulsatile cell behaviors observed in optogenetic studies using the RhoGEF2 DHPH catalytic domains in the early embryo (16). The more heterogenous cell area reductions and absence of tissue folding that we observe in the germband of optoGEF embryos, compared to in prior optogenetic studies conducted at earlier stages in development, may reflect a number of factors: differing gene expression patterns at these developmental stages, existence of anisotropies associated with planar polarity in the germband, size of the optogenetic activation region, mechanical properties of the germband tissue, mechanical or geometrical constraints, or differences in the optogenetic tools themselves.

During axis elongation, myosin II is enriched at the vertical AP cell junctions and in the medial-apical domain, producing a planar polarized pattern of myosin (22–24, 29–33) that is thought to produce anisotropic contractile forces that drive cell intercalation and convergent extension (22, 23, 29, 30, 32, 33, 35). The effects of optoGEF and optoGAP perturbations on myosin planar polarity in the germband are rapid. During the first 2.5 min, myosin planar polarity reduction occurs faster in optoGAP than in optoGEF embryos, but optoGEF embryos show a greater final reduction in planar polarity than optoGAP. These distinct dynamics may reflect the molecular mechanisms and timescales for Rho1 activation versus inactivation, Rho-kinase activation versus inactivation, and myosin regulatory light chain phosphorylation versus dephosphorylation. In addition, Rho1 and RhoGEF2 are known to play roles in regulating the actin cytoskeleton (68), which could contribute to the observed changes in myosin localization patterns and cell behaviors. Interestingly, while myosin planar polarity was more strongly reduced in optoGEF compared to optoGAP embryos, we observed higher rates of cell rearrangements in the optoGEF case, reflecting that other aspects of the myosin localization pattern may influence contractile cell behaviors that contribute to axis elongation. One possibility is that the enhanced medial myosin and pulsatile apical cell area fluctuations in optoGEF embryos may promote cell rearrangement and intercalation (discussed below).

The medial-apical myosin pattern in optoGEF embryos was similar to that observed in the germband of embryos overexpressing RhoGEF2, expressing constitutively active concertina, or treated to depolymerize microtubules (48, 50, 61). Interestingly, the medial myosin pattern in optoGEF embryos was distinct from the more junctional myosin pattern in embryos expressing a phosphomimetic myosin regulatory light chain locked-in to the “on” state (30), possibly suggesting additional effects of the optoGEF tool on the actomyosin meshwork beyond increasing myosin activity. Alternatively, the difference in myosin localization between optoGEF and phosphomimetic embryos might reflect depletion of myosin from junctions due to a limited myosin pool or changes in the balance between membrane localization of RhoGEF2 and Cysts/Dp114RhoGEF, which are thought to have distinct roles in regulating medial and junctional myosin in the germband (49, 50). The condensed cluster of medial myosin observed in optoGEF embryos is reminiscent of the myosin in apically constricting cells under isotropic tension in the invaginating posterior midgut of wild-type embryos and in the presumptive mesoderm of embryos with disrupted tension anisotropy (62, 69), which raises the possibility that the optogenetically-induced change in myosin pattern alters the tension distribution in the tissue. Furthermore, myosin recruitment to the cell cortex can be mechanosensitive (33), raising the possibility of complex effects of the optogenetic perturbations both within and outside of the germband. Future studies to quantify the mechanical tension in the tissue will be needed to directly link changes in myosin localization with changes in mechanical tension in the germband.

During axis elongation, accumulation of actomyosin at the medial-apical domain of germband cells is associated with fluctuations in apical cell area in wild-type embryos (37, 66). We found that the optogenetic perturbations tuned the magnitude of these pulsatile cell area changes. In optoGEF embryos, the accumulation of myosin in the medial-apical domain and formation of a radial myosin pattern was associated with increased magnitude of area fluctuations, enhanced net area changes, and high cell area heterogeneity compared to in control embryos. The increase in cell area heterogeneity in the germband contrasts with the more uniform cell area decreases following optogenetic activation of Rho1 in localized regions of tissue in the early embryo (16, 20). One possible explanation is that a tissue-wide pattern of medial myosin accumulation has different effects on apical cell area than a local pattern, potentially due to the size of the contractile region and the geometrical and mechanical constraints in the embryo. Alternatively, aspects of the endogenous planar polarity pattern in the tissue may contribute to this phenotype. Future studies that locally activate Rho1 signaling in the germband will be essential to understanding the effects of local versus global activation in a planar polarized tissue.

While a population of cells in optoGEF embryos displayed progressive cell area decreases, other cells displayed a combination of transient periods of area reduction and expansion with no stabilization of cell area changes, similar to in *bcd nos tsl* mutant embryos that have high medial relative to junctional myosin levels (37). The absence of a net reduction in apical cell area over time despite increased medial myosin might be caused by the concomitant reduction of junctional myosin in these embryos, which could disrupt tension stabilization among groups of cells. Consistent with this notion, progressive apical constriction was observed in ventrolateral germband cells in JAK/STAT mutant embryos, which have high levels of medial-apical myosin but normal levels of junctional myosin (36). A role for junctional myosin in stabilization of cell area was also evident in optoGAP embryos, where the reduction in junctional and medial myosin generated a subtle area relaxation in a considerable fraction of cells. In addition, the observed changes in myosin localization in optoGEF and optoGAP embryos could be coupled with changes in the actin cytoskeleton or cell adhesions, which might contribute to these phenotypes. Future studies analyzing the effects of these optogenetic perturbations on other cytosketetal and adhesive proteins will be needed to distinguish between these possibilities.

Interestingly, the amplitudes of apical cell area fluctuations in optoGEF and optoGAP embryos, in which myosin planar polarity is disrupted, correlate with the rates of cell rearrangement in the germband. As myosin planar polarity is thought to generate the anisotropic internal tensions that drive cell rearrangement, a complete lack of myosin planar polarity would be predicted to block cell rearrangements in the germband. While myosin planar polarity is nearly abolished in both optoGEF and optoGAP embryos, cell rearrangements are less severely impacted in optoGEF embryos. One explanation is that small remaining differences in myosin localization and activity at junctions is not detectable in imaging but still sufficient to drive rearrangements. Alternatively, mechanisms that are independent of junctional myosin planar polarity might be contributing to cell rearrangement in this context (70). One potential mechanism is that active myosin-associated cell area fluctuations might help promote cell rearrangements. Consistent with this notion, recent theoretical and experimental work highlights that cells must overcome a physical energy barrier associated with cell shape changes in order to proceed through a cell rearrangement (43–45). Active fluctuations in tension and shape can help to overcome these energy barriers and promote the ability of a tissue to remodel and flow—tissue fluidity. Indeed, recent experimental studies point toward exactly such a role for active fluctuations in promoting fluidity within tissues (40, 41). Future studies of the biophysics of cell rearrangements in the germband will be needed to explore further how active fluctuations might contribute to oriented cell rearrangements in both wild-type and mutant embryos.

The optogenetic perturbations to Rho1 activity in the germband epithelium in this study allowed us to link cellular junctional and medial myosin patterns to distinct aspects of cell behavior that contribute to convergent extension. However, the rapid changes in the myosin pattern in the germband of optoGEF embryos—from the endogenous planar polarized pattern to one reminiscent of the radial patterns present in invaginating cells of the presumptive mesoderm— were not sufficient to completely transform cell behaviors to recapitulate coordinated apical constriction and tissue invagination. This points to the existence of additional mechanisms beyond myosin pattern that are required to ectopically recapitulate morphogenetic movements at other stages of development. Going forward, it will be interesting to further explore how internal biochemical regulation couples with mechanical cues and constraints to give rise to the dynamic myosin patterns that orchestrate diverse cell behaviors and morphogenetic events. In addition, it will be important to investigate how anisotropic patterns of junctional actomyosin couple with active fluctuations driven by the apical actomyosin cytoskeleton to promote cell rearrangements and epithelial tissue fluidity during development. The optogenetic tools and approaches we developed here will be valuable in these investigations.

## AUTHOR CONTRUBUTIONS

R.M.H. designed research, performed research, analyzed data, and wrote manuscript; C.C. analyzed data; C.A. analyzed data; A.L. analyzed data; K.E.K. designed research, analyzed data, and wrote manuscript.

## ACKNOWLEDGMENTS

We thank Stefano De Renzis for the UASp>CIBN::pmGFP fly stock, Jennifer Zallen and Frederik Wirtz-Peitz for the UASp-attB destination vectors, the Bloomington Drosophila Stock Center (BDSC) for fly stocks, BACPAC Genomics at Children’s Hospital Oakland Research Institute (now BACPAC Genomics) for BAC clones, and members of the Kasza Lab for helpful discussion and comments on the manuscript. This work was supported by NSF Civil, Mechanical, and Manufacturing Innovation Grant 1751841 (to K.E.K.). K.E.K holds a Career Award at the Scientific Interface from the Burroughs Wellcome Fund, a Clare Boothe Luce Professorship, and a Packard Fellowship.

## MATERIALS AND METHODS

### Cloning

To generate CRY2-RhoGEF2, the full RhoGEF2 cDNA of *Drosophila melanogaster* was amplified by PCR from plasmid UAS-akkordion dNhe (Addgene 41976) (71), and the cDNA of CRY2PHR was PCR amplified from the pCRY2PHR-mCherryN1 plasmid from the Tucker Lab (Addgene 26866) (7). DNA fragments were linked using the NEB Builder HiFi DNA Assembly Kit (New England BioLabs). RhoGAP71E was PCR amplified from the exons corresponding to RhoGAP71E protein from pACMAN CH321-72J07 (BACPAC resources), and DNA fragments were assembled using a combination of restriction-ligation and DNA fragment assembly to obtain the full RhoGAP71E cDNA. To produce an N-terminal fluorescently-tagged version of constructs, the sequence encoding mCherry from the pCRY2PHR-mCherryN1 plasmid from the Tucker Lab (Addgene 26866) (7) was included during DNA fragment assembly. In the constructs, a (GA)5 linker was introduced between mCherry and CRYPHR and a GG(SG)4 linker was introduced between CRY2 and RhoGEF2 or RhoGAP71E. Constructs were cloned into the pENTR/D-Topo vector (Life Technologies) and recombined into UASp-attB destination vector (gift of F. Wirtz-Peitz) using the Gateway cloning system (Life Technologies) for subsequent expression using the Gal4/UAS system (72). The resulting plasmids were purified, sequenced, and used to generate transgenic flies (Bestgene). To ensure comparable expression levels, all transgenes corresponding to CRY2 constructs were inserted into the attP2 site on chromosome III.

### Fly stocks

Stocks and crosses were maintained at 23°C and experiments were performed at room temperature (∼ 21°C). The stocks w*;; w+, UASp>mCherry-CRY2PHR-RhoGEF2 and w*;; w+, UASp>mCherry-CRY2PHR-RhoGAP71E were generated in this study, crossed with w*; P[w+, UASp>CIBNpm-GFP]/Cyo; Sb/TM3,Ser (gift of Stefano De Renzis, EMBL, Heidelberg, Germany), and subsequently expressed using the maternal α -tubulin matα -tub15 or matα -tub67 Gal4-VP16 drivers (mat67, mat15, gift of D. St Johnston). To visualize myosin II, mat15 and sqh>sqh-mCherry (BDSC 59024, donated by Beth Stronach, University of Pittsburgh) were recombined on chromosome III and used alone or in combination with mat67. Crosses and embryos were kept protected from light. Fly sorting was performed in the dark on a stereomicroscope equipped with a Red 25 Wratten Filter. Embryos studied were progeny of females of the following genotypes:

UASp>CIBNpm-GFP/ mat67 (II); sqh>sqh-mCherry/+ (III)

UASp>CIBNpm-GFP/+ (II); UASp>mCherry-CRY2PHR-RhoGEF2/ mat15 (III)

UASp>CIBNpm-GFP/+ (II); UASp>CRY2PHR-RhoGEF2/ sqh>sqh-mCherry, mat15 (III)

UASp>CIBNpm-GFP/ mat67 (II); UASp>mCherry-CRY2PHR-RhoGAP71E/ mat15 (III)

UASp>CIBNpm-GFP/ mat67 (II); UASp>CRY2PHR-RhoGAP71E / sqh>sqh-mCherry, mat15 (III)

Fly stocks used in this study:

w*;; w+, UASp>mCherry-CRY2PHR-RhoGEF2/TM3,Sb (this study) w*;; w+, UASp>CRY2PHR-RhoGEF2/TM3,Sb (this study)

w*;; w+, UASp>mCherry-CRY2PHR-RhoGAP71E/TM3,Sb (this study) w*;; w+, UASp>CRY2PHR-RhoGAP71E/TM3,Sb (this study)

w*; w+, UASp>CIBNpm-GFP/Cyo; Sb/TM3,Ser (Stefano De Renzis, EMBL, Heidelberg, Germany)

w*;; sqh>sqh-mCherry (Beth Stronach, University of Pittsburgh)

w*; w+, UASp>CIBNpm-GFP, mat67; sqh>sqh-mCherry

w*;; sqh>sqh-mCherry, mat15

w*; mat67; sqh>sqh-mCherry, mat15

### Assessing axis elongation phenotypes and embryo viability

Samples for imaging were prepared in a dark room illuminated with red light. Embryos were collected under red light, maintained in darkness for 4 h after being laid, briefly exposed to red light to manually score body axis elongation phenotypes on a stereomicroscope, and returned to darkness. After one day, embryo viability was assessed from the embryo hatching rate.

### Live imaging

Samples for imaging were prepared in a dark room illuminated with red light. Embryos in early stage 6 were selected under halocarbon oil 27 (Sigma), dechorionated with 50% bleach for 2 min, washed with water, and mounted using a mixture 50:50 of halocarbon oil 27:700 on a custom-made imaging chamber between an oxygen-permeable membrane (YSI Incorporated) and a glass coverslip. Embryos were positioned ventrolaterally for observation of the germband and were imaged on a Zeiss LSM 880 confocal microscope equipped with a diode laser for 561 nm excitation and an Argon laser for 488 excitation, a standard LSM confocal detector, and an Airyscan detector with FAST module (Carl Zeiss, Germany). Embryos were photoactivated with 488 nm light (Carl Zeiss, Germany). Imaging was performed with C-Apo 40X/1.2 NA water immersion objective or a Plan-Apo 63X/1.40 NA oil immersion objective (Carl Zeiss, Germany). Images were acquired using the standard LSM mode unless otherwise noted. For myosin imaging, the Airyscan FAST module was utilized. The light for bright field illumination was filtered through a Red 25 Kodak Wratten Filter to prevent unwanted photoactivation during embryo selection. Image acquisition was performed with ZEN Black software. Images of the germband were acquired as 10 μm z-stacks with a 1 μm z-step (40X objective, LSM mode) or a 0.7 μm z-step (63X objective, Airyscan FAST), beginning at the apical side of the tissue. Z-stacks were acquired every 15 s. Photoactivation was achieved by scanning the same ventrolateral region of the germband with blue laser light (λ = 488 nm) that was being imaged with blue and/or green light (λ = 488 nm, λ = 561 nm), at every imaging z-plane and time-step. To visualize myosin prior to photoactivation, mCherry-tagged myosin regulatory light chain was imaged with 561 nm light for 2.5 min. For quantification of protein localization patterns at different positions along the z-axis, a 10 µm or 15 µm z-stack (beginning at the apical side) with a 0.7 µm or 0.5 μm z-step was taken prior to and after photoactivation; photoactivation was achieved by scanning the blue laser (λ = 488 nm) over a 10 μm z-stack with 1 μm z-steps every 15 s for 2.5 min (40X objective, LSM mode).

### Image analysis and quantification

Tissue elongation during body axis elongation was calculated from confocal time-lapse movies using the particle image velocimetry (PIV) software PIVlab version 1.41 (73) in MATLAB as previously described (30, 73). For myosin analysis, still frames shown and confocal movies analyzed correspond to the maximum intensity projections of 5 μm at the apical side of the tissue, starting apically just underneath the vitelline membrane, unless otherwise noted. Images were processed and analyzed using the ImageJ distribution Fiji (74, 75). The mean junctional myosin intensity at each cell edge was quantified manually using 0.5 μm-wide lines (excluding vertex regions) in a 40 μm square region, and the intensity and orientation of each edge was obtained. Medial-apical myosin intensity was quantified manually as the mean intensity in a polygon manually drawn just inside the cell membrane outline. At least 10 cells and 40 junctions were analyzed per embryo. Planar polarity was obtained as the ratio of the mean intensity of AP (“vertical”) edges to the mean intensity of DV (“horizontal”) edges for each embryo. Cell outlines were visualized using CIBNpm-GFP for manually obtaining cell areas. At least 6 cells per embryo and 3-4 embryos per conditions were analyzed. For quantifying protein localization patterns at different z-positions along the apical to basal axis, 0.5 μm-wide lines (excluding vertex regions) were used to measure intensity at cell edges and polygons drawn just inside the cell outline were used to quantify medial/cytoplasmic intensity. For automated image segmentation analysis, the SEGGA tissue segmentation and analysis software was used, cell outlines were visualized with CIBNpm-GFP, and 3-5 embryos per genotype were analyzed. (Figures 1*C* and 3*B,D*) (24).

### Cell behavior analysis

Cell horizontal length, coefficient of variation of apical cell area, cell rearrangements, and tissue elongation were calculated from cell segmentation analysis using the SEGGA software (24), and 3-5 embryos were analyzed per genotype. Analyses of single cell areas, medial myosin, and junctional myosin were performed manually. 8-12 cells per embryo and 3-4 embryos per genotype were analyzed. For Figures 4 and 5, raw cell area and myosin intensity data were treated with a Savitzky-Golay filter (filter window: 9, order: 5). Myosin intensities were normalized to the value at *t* = 0. Changes of cell area and myosin correspond to the differences between 15 s time steps.

### Statistical Analysis

Mean values and standard error of the mean (SEM) were determined as indicated in each figure legend. The non-parametric Kruskal-Wallis test with post-hoc Dunn test was used to determine p-values unless otherwise noted.

## Notes

### Competing Interest Statement

The authors have declared no competing interest.

